# Multiple sex chromosomes of *Yponomeuta* ermine moths suggest a role of sexual antagonism in sex chromosome turnover in Lepidoptera

**DOI:** 10.1101/2023.06.06.543653

**Authors:** Provazníková Irena, Dalíková Martina, Voleníková Anna, Roessingh Peter, Sahara Ken, Provazník Jan, Marec František, Nguyen Petr

## Abstract

Sex chromosome-autosome fusions give rise to neo-sex chromosomes, which provide an insight into early evolution of sex chromosomes and drivers of chromosomal fusions. While sex chromosome-autosome fusions are scarce in vertebrates with female heterogamety (♀ZW/♂ZZ), they are common in moths and butterflies (Lepidoptera), the most species rich group with heterogametic females. This contradicts theoretical model that assumes chromosome fusions to be random and predicts them to be rare in taxa with high chromosome number such as Lepidoptera. In the present study we analyzed sex chromosomes in nine ermine moths of the genus *Yponomeuta* (Yponomeutidae) and their two outgroups, *Teinoptila gutella* (Yponomeutidae) and *Plutella xylostella* (Plutellidae). We employed genomic *in situ* hybridization to identify sex chromosomes and used a custom designed microarray to identify Z-linked genes. Our results confirmed a multiple sex chromosome system Z1Z2W to be present in *T. gutella* and all *Yponomeuta* spp. except for *Y. tokyonella*. The multiple sex chromosome system resulted from a fusion between the W chromosome and autosome homeologous to the *Bombyx mori* chromosome 2 (BmChr2). The BmChr2 bears a cluster of genes with ovary-specific expression which suggests that sexually antagonistic selection could have driven fixation of the fusion in a common ancestor of *Yponomeuta* and *Teinoptila* genera. We hypothesize that sex chromosome turnover in Lepidoptera could be driven by sexual antagonism.

## Introduction

Sex chromosomes represent a specific part of genome. They are subject to a selection regime distinct from autosomes and play an important role in evolution (Payseur et al. 2018; Connallon et al. 2018). The canonical model of sex chromosome evolution postulates that sex chromosomes arise from a pair of autosomes. When this pair acquires a sex determining (SD) locus, one of its alleles is limited to a heterogametic sex (XY males or ZW females). Selection should favor a linkage disequilibrium between the sex-limited allele and sexually antagonistic (SA) mutations, i.e. mutations beneficial to the heterogametic but detrimental to the homogametic sex (XX females or ZZ males). Suppression of recombination between SD and SA loci is then advantageous and results in differentiation of the sex chromosome pair via accumulation of repeats and deleterious mutations due to Hill-Robertson interactions (Wright et al. 2016; Kratochvíl et al. 2021).

The SA selection should also drive evolution of neo-sex chromosomes resulting from sex chromosome-autosome fusions (Charlesworth and Charlesworth, 1980; Kitano et al., 2009; but see Pennell et al., 2018; Anderson et al., 2020), which could provide an insight into sex chromosome evolution in taxa with old and highly differentiated sex chromosome systems such as insects or vertebrates (Blackmon et al. 2017; Stöck et al. 2021). It has been proposed that sex chromosome-autosome fusions are rare in vertebrate taxa with female heterogamety (Pokorná et al. 2014; Pennell et al. 2018). However, steadily growing number of neo-sex chromosomes have been reported in moth and butterflies (Lepidoptera), which comprise the most speciose group with female heterogamety (Nguyen and Carabajal Paladino 2016; Carabajal Paladino et al. 2019; Smith et al. 2019; Yoshido et al. 2020). Moreover, neo-sex chromosomes have been proposed to play a role in adaptive evolution and diversification of Lepidoptera (Nguyen et al. 2013; Smith et al. 2019; Yoshido et al. 2020). Sex chromosome-autosome fusions increase a number of sex-linked genes, which could significantly accelerate the accumulation of genetic incompatibilities between populations (Turelli and Begun 1997). Autosomes fused with sex chromosomes in large lepidopteran radiations were enriched for clusters of genes involved in detoxification and regulated absorption of plant secondary metabolites, which are crucial for larval performance on their host plants (Nguyen et al. 2013; Carabajal Paladino et al. 2019). Physical linkage between performance and sex-linked genes for either female host preference (Thompson 1988; Nygren et al. 2006) or host-independent reproductive isolation (Sperling 1994; Presgraves 2002) could facilitate adaptation and speciation in the presence of gene flow (Matsubayashi et al. 2010).

As recombination ceases due to achiasmatic meiosis in lepidopteran females, maternally transmitted (neo-W-linked) alleles of performance genes deteriorate which could contribute to population-specific divergence (Filatov 2018). Furthermore, selection for dosage compensation caused by increasing environmental stress, is then expected and beneficial duplications of performance genes can be fixed by positive selection (Innan and Kondrashov 2010; Singh et al. 2020). Such amplification and functional divergence of performance genes upon sex chromosome-autosome fusions was proposed to be a key innovation, which enhanced adaptive radiation of leaf-rollers of superfamilies Tortricoidea and Gelechioidea (Nguyen et al. 2013; Carabajal Paladino et al. 2019).

Small ermine moths of the genus *Yponomeuta* (Yponomeutoidea, Yponomeutidae) have been a subject of multidisciplinary investigation of the evolution of insect-plant associations and speciation of phytophagous insects (Menken et al. 1992; Menken 1996; Menken and Roessingh 1998), and pheromone communication (Löfstedt et al. 1991; Liénard and Löfsted 2010). All ermine moths but one are monophagous and have an ancestral association with host plants of the family Celastraceae (Menken 1996). However, during their dispersion from East Asia to the western Palearctic, the common ancestor of the European clade switched its hosts from the family Celastraceae to Rosaceae and Salicaceae (Turner et al. 2010).

Analysis of meiotic nuclei of interspecific hybrids of the European ermine moths *Yponomeuta cagnagella* and *Y. padella* revealed numerous pairing irregularities including characteristic loops between paired homologues in meiosis, which indicate chromosome inversions (Hora et al. 2019). Saitoh (1960) reported male chromosome numbers of four *Yponomeuta* species. The karyotypes of *Y. malinella* and *Y. sedella* males corresponded to the ancestral lepidopteran karyotype n=31 exemplified by yponomeutoids *Atteva aurea* (Attevidae), *Zelleria haimbachi* (Yponomeutidae), and *Plutella xylostella* (Plutellidae) (Ennis, 1976; Kawazoé, 1987; cf. Van’t Hof et al., 2013). By contrast, the chromosome number of *Y. polystictus* and *Y. sociatus* males was only n=30. The difference may be due to a sex chromosome-autosome fusion identified in females by Nilsson et al. (1988), who found a derived sex chromosome system Z1Z2W (2n♀=61, 2n♂=62) in six ermine moths of the *Y. cagnagellus*–*irrorellus* clade. The *Yponomeuta* ermine moths thus represent an ideal system for study a role of neo-sex chromosomes in ecological adaptation and speciation of Lepidoptera.

In the present study, we employed genomic *in situ* hybridization to identify sex chromosomes in 9 *Yponomeuta* spp. and their two outgroups, *Teinoptila gutella* (Yponomeutidae) and *Plutella xylostella* (Plutellidae). In *Y. evonymella*, we built genomic resources such as transcriptome sequence and a genomic library of bacterial artificial chromosomes (BACs) and used a custom designed microarray to identify Z-linked genes. Quantitative PCR (qPCR) with male and female genomic DNA (gDNA) and physical mapping by means of fluorescence *in situ* hybridization with BAC-derived probes confirmed that the *Y. evonymella* Z2 chromosome corresponds to an autosome homeologous to the *Bombyx mori* chromosome 2 (BmChr2). The BmChr2 bears a cluster of genes with ovary-specific expression which suggests that sexually antagonistic selection could have driven fixation of sex chromosome-autosome fusion in a common ancestor of *Yponomeuta* and *Teinoptila* spp.

## Material and methods

### Insects

All examined species were collected from wild populations except for *P. xylostella* which was obtain from a laboratory stock. The specimens were mostly collected as larvae and either processed immediately upon collection or reared under ambient conditions on their host plants (for details see **Suppl. Table S1**).

### Chromosomal preparation

Meiotic and mitotic chromosomes were obtained from female and male gonads of 5th instar larvae by spreading technique as described in (Provazníková et al. 2021). Chromosomal preparations were afterwards dehydrated in ethanol series (70%, 80% and 100%, 30 sec each) and stored at -20°C or - 80°C until further use.

### RNA sequencing and transcriptome assembly

Total RNA was extracted from the *Y. evonymella* female larva with its gut removed using an RNA Blue reagent (Top-Bio, Prague, Czech Republic) following the manufacturer’s protocol. The Illumina mRNA-seq library was constructed and sequenced on the Illumina HiSeq2000 platform by EMBL Genomics Core Facility (Heidelberg, Germany). Resulting raw 100-bp paired-end reads were trimmed and quality filtered by Trimmomatic version 0.30 (‘LEADING:5 TRAILING:5 SLIDINGWINDOW:4:20’; Bolger et al., 2014). Transcriptome sequence was then assembled *de novo* by SOAPdenovo-trans-127mer (Xie et al. 2014) with multiple k-mer sizes ranging from 35 to 75 in increments of 10 and Trinity with the ‘--SS_lib_type RF’ option (Haas et al. 2013). The resulting assemblies were merged and redundancy was removed using the EvidentialGene pipeline (Gilbert 2013). The raw reads were deposited in NCBI under SRA accession number PRJNA788289.

### Array-CGH analysis

To identify sex-linked genes in *Y. evonymella*, we performed comparative genomic hybridization on a microarray (array-CGH) following (Baker and Wilkinson 2010). We searched for 1:1 orthologs of *B. mori* genes using the EvidentialGene dataset (see above) as input for HaMStR (Ebersberger et al. 2009) with the ‘-representative’ option and lepidopteran core ortholog set by (Breinholt and Kawahara 2013). The *Y. evonymella* orthologs were used for design of 60-mer oligonucleotide probes for a custom-made microarray slide using Agilent Technologies eArray design wizard (https://earray.chem.agilent.com/earray/). Female and male gDNA were extracted from larvae using CTAB protocol by (Winnepenninckx et al. 1993). DNA was quantified using the Qubit dsDNA BR Assay kit (Thermo Fisher Scientific, Waltham, MA, USA). DNA digestion, labelling, and array-CGH were performed by GenLabs (Prague, Czech Republic) according to a protocol for Agilent oligonucleotide array-based CGH for gDNA analysis. Hybridization intensities were extracted using Agilent’s Feature Extraction software. Filtering and analysis of feature intensities followed (Baker and Wilkinson 2010) implemented in the custom Python script (Yoshido et al. 2020). The cut-off value of 0.5 was used to identify Z-linked genes.

### *Y. evonymella* BAC library and screening

Genomic library of BACs was constructed for *Y. evonymella* by AC Amplicon Express (Pullman, WA, USA). High molecular weight gDNA of *Y. evonymella* males was partially digested by *Hind* III and cloned into the pCC1BAC (Epicentre, Madison, WI, USA) vector, which was transformed into the DH10B *Escherichia coli* cells. The library consists of 20 736 clones with the average insert of about 125 Kbp. To identify BAC clones bearing genes of interest, i.e. a particular plate, row, and column of the BAC library, the library was screened by means of PCR according to the manufacturer’s guidebook. For the screening procedure, individual BAC clones were pooled into 18 sets (or superpools) by the manufacturer, each containing 1 152 BAC clones. First, screening of superpools is performed to identify which contain BAC clone(s) with the sequence of interest. Each superpool corresponds to another subset of BACs comprising 21 matrixpools screened by second PCR. The matrixpools are plate, row, and column pools combined in a way, which allows identification of the clone bearing the gene of interest.

The 10µl PCR reaction contained 1x reaction buffer, 3 µM of each primer (**Suppl. Table S2)**, 1 µl of template gDNA, 0.2 mM dNTPs and 0.5 U TaKaRa rTaq DNA polymerase (TaKaRa, Otsu, Japan). The amplification was carried out by PCR involving a denaturation step at 94 °C for 3 min; followed by 30 cycles of a denaturation at 94 °C for 30 s, an annealing at 58-60 °C for 45 s and an elongation at 72 °C for 45-60 s; and final elongation at 72 °C for 3 min. Afterwards, PCR reactions were visualized by gel electrophoresis and evaluated according to manufacturer’s manual. To get single colonies, the positive BAC clones were plated on agar plate containing chloramphenicol (25 µg/ml). Presence of desired sequence was again verified by PCR on several single colonies per plate. PCR was prepared and carried out as described above using the single colonies as a template.

### *In situ* hybridization

Genomic DNA for fluorescence *in situ* hybridization (FISH) experiments was extracted from male and female larvae or pupae by standard phenol-chloroform method (Blin and Staford 1976). Obtained gDNA was amplified using illustra GenomiPhi HY DNA Amplification Kit (GE Healthcare, Milwaukee, WI, USA) according to manufacturer’s protocol, purified by precipitation with isopropanol and sodium acetate, and dissolved in ultra clean water. DNA from selected BACs for was extracted by Plasmid Midi Kit (QIAGEN, Hilden, Germany) according to manufacturer’s protocol.

Telomeric probe, female gDNA and BAC DNA were labelled by the nick translation using nick translation kit (Abbott Molecular Inc., Des Plaines, IL, USA) or a protocol described in (Kato et al. 2006) with some modifications (Dalíková et al. 2017). Using the nick translation kit (Abbott Molecular Inc.), the 25µl labelling reaction contained 500 ng DNA, 40 μM dATP, 40 μM dCTP, 40 μM dGTP, 14.4 μM dTTP and 25.6 μM Cy3-dUTP (Jena Bioscience, Jena, Germany) for telomeric probe; fluorescein-12-dUTP (Jena Biosciences) for female gDNA; Cy3-dUTP and fluorescein-12-dUTP (both Jena Biosciences) for BAC DNA. The reaction was incubated at 15°C for 75 min for telomeric probe, 4 h for female gDNA and 5 h for BAC DNA. The modified (Kato et al. 2006) nick translation reaction contained 1 µg of unlabelled DNA; 0.5 mM dATP, dCTP and dGTP; 0.1 mM dTTP; 20 µM of labelled nucleotides; 1x nick translation buffer (50 mM Tris-HCl pH 7.5, 5 mM MgCl2, 0.005% BSA), 10 mM ?-mercaptoethanol, 2.5 × 10-4 U DNase I (ThermoFisher Scientific, Waltham, MA USA) and 1 U DNA polymerase I (ThermoFisher Scientific). The reaction was incubated at 15°C for 60 min to label telomeric probe, 210 min for female gDNA and 5 h for BAC DNA.

Genomic *in situ* hybridization (GISH) combined with telomeric probe and FISH with bacterial artificial chromosome (BAC-FISH) were carried out as described in (Yoshido et al., 2005a) and (Yoshido et al., 2005b), respectively, with some modifications. Briefly, the hybridization cocktail contained labelled probes, 500 ng fluorescein-labelled female gDNA and 100 ng of Cy3-labelled telomeric or 300 ng Cy3-labelled BAC DNA and 500 ng fluorescein-labelled BAC DNA, 3 µg of male competitor gDNA fragmented by heat for 20 min at 99°C, 25 μg sonicated salmon sperm DNA (Sigma-Aldrich, St. Louis, MO, USA) in 10 µl of 50% deionized formamide and 10% dextran sulfate in 2x SSC. The hybridization mixture was denatured for 5 min at 90°C. Chromosome slides were denatured in 70% formamide in 2x SSC for 3.5 min at 68°C. After 3-days hybridization at 37°C, slides were washed for 5 min in 0.1x SSC with 1% Triton X-100 at 62°C and counterstained with 0.5 μg/ml DAPI (4’,6-diamidino-2-phenylindole; Sigma-Aldrich) in antifade with DABCO (1,4-diazabicyclo (2.2.2)-octane; Sigma-Aldrich).

### Reprobing

To physically map multiple BACs, two rounds of BAC-FISH on the same chromosome slides were carried out due to the limiting number of available fluorochromes. The reprobing procedure was done as described in Zrzavá et al. (2018) with some modifications. The slides were incubated in 2x SSC for 30 min to remove cover slips and incubated for 10 min in 50% formamide, 1% Triton X in 0.1x SSC at 70°C to denature and eliminate the first probes. Afterwards, the slides were placed into prechilled 70% ethanol for 1 minute and then dehydrated at room temperature in 80% and 100% ethanol (30 s each). When dried, the slides were immediately used for another hybridization with BAC probes as described above.

### Documentation and image processing

Preparations from FISH experiments were observed in a Zeiss Axioplan 2 microscope (Carl Zeiss, Jena, Germany) with a fluorescence filter sets and a monochrome CCD camera XM10 (Olympus Europa Holding, Hamburg, Germany). The images were captured in black-and-white, separately for each fluorescent dye with cellSens Standard software version 1.9 (Olympus). Subsequently, the images were pseudocolored and merged in Adobe Photoshop CS4 (version 11).

### Quantitative PCR

To confirm results from array-CGH and BAC-FISH, and to verify a common origin of the Z2 chromosome across the genus *Yponomeuta*, testing for sex-linkage of selected genes by quantitative PCR (qPCR) was performed in selected species, namely *Y. plumbella, Y. evonymella* and *Y. tokyonella*. Experiment was designed and carried out according to Nguyen et al. (2013) with some modifications. The qPCR analyses were performed with *Acetylcholinesterase 2 (Ace2)* as an autosomal reference, genes *Henna* and *Kettin* as markers for the ancestral Z1 chromosome and genes *Arp6* and *Plep1* as markers for the Z2 chromosome. The reference gene and genes of interest were analysed simultaneously in three biological and technical replicas for both males and females. For qPCR experiments, gDNA was extracted from male and female individuals of larval or pupal stage by NucleoSpin Tissue kit (Macherey-Nagel, Düren, Germany) or NucleoSpin Tissue XS kit (Macherey-Nagel) according to manufacturer’s protocol. One 10µl reaction contained 1-10 ng of gDNA, 0.4 or 0.8 µM each primer (details in **Suppl. Tab. S3**) and 5 µL of SYBR Mix (Xceed qPCR SG 2x Mix Lo-ROX, IAB, Prague, Czech Republic). The experiment was carried out using the C1000 Thermal cycler CFX96 Real-Time System (Bio-Rad, Hercules, CA, USA). Data were analysed using software Bio-Rad CFX Manager 3.1. To determine the amplification efficiency of the reaction for each gene (*E*); 0x, 5x, 25x and 125x dilution series of gDNA pool from all gDNA samples of each species was analysed. Using the formula *R* = [(1+ *E*Reference) CtReference] / [(1+ *E*Target)CtTarget] (Rovatsos et al. 2014), the target to reference gene dose ratio (*R*) was calculated for each biological sample. The statistical analysis was carried out as described in Dalíková et al. (2017). Briefly, two hypothesis, autosomal hypothesis (*R* value ratio male:female to be 1:1) and Z-linkage hypothesis (*R* value ratio male:female to be 2:1) were tested by unpaired two-tailed t test for unequal variances.

## Results

## Karyotype analysis

Diploid chromosome numbers in *P. xylostella* (2n=62), *Y. cagnagella* (2n=♀61/♂62), *Y. padella* (2n=♀61/♂62) and *Y. evonymella* (2n=♀61/♂62) were already known from previous studies (Kawazoé 1987; Nilsson et al. 1988) and verified in this study. Based on mitotic metaphases and meiotic bivalents, a diploid chromosome number was determined to be 2n=♀61/♂62 in remaining species, namely *T. gutella, Y. plumbella, Y. polysticta, Y. kanaiella* and *Y. mahalebella* **(Fig. 1)** with exception of *Y. tokyonella* where reduced chromosome number was observed (2n=60) **(Fig. 2d)**. In *Y. orientalis*, a diploid chromosome number was also estimated to be 2n=♀61/♂62, however, due to lack of mitotic chromosome preparations, more nuclei and specimen need to be examined to verify the preliminary results **(Fig. 1i, j)**.

**Figure 1:**
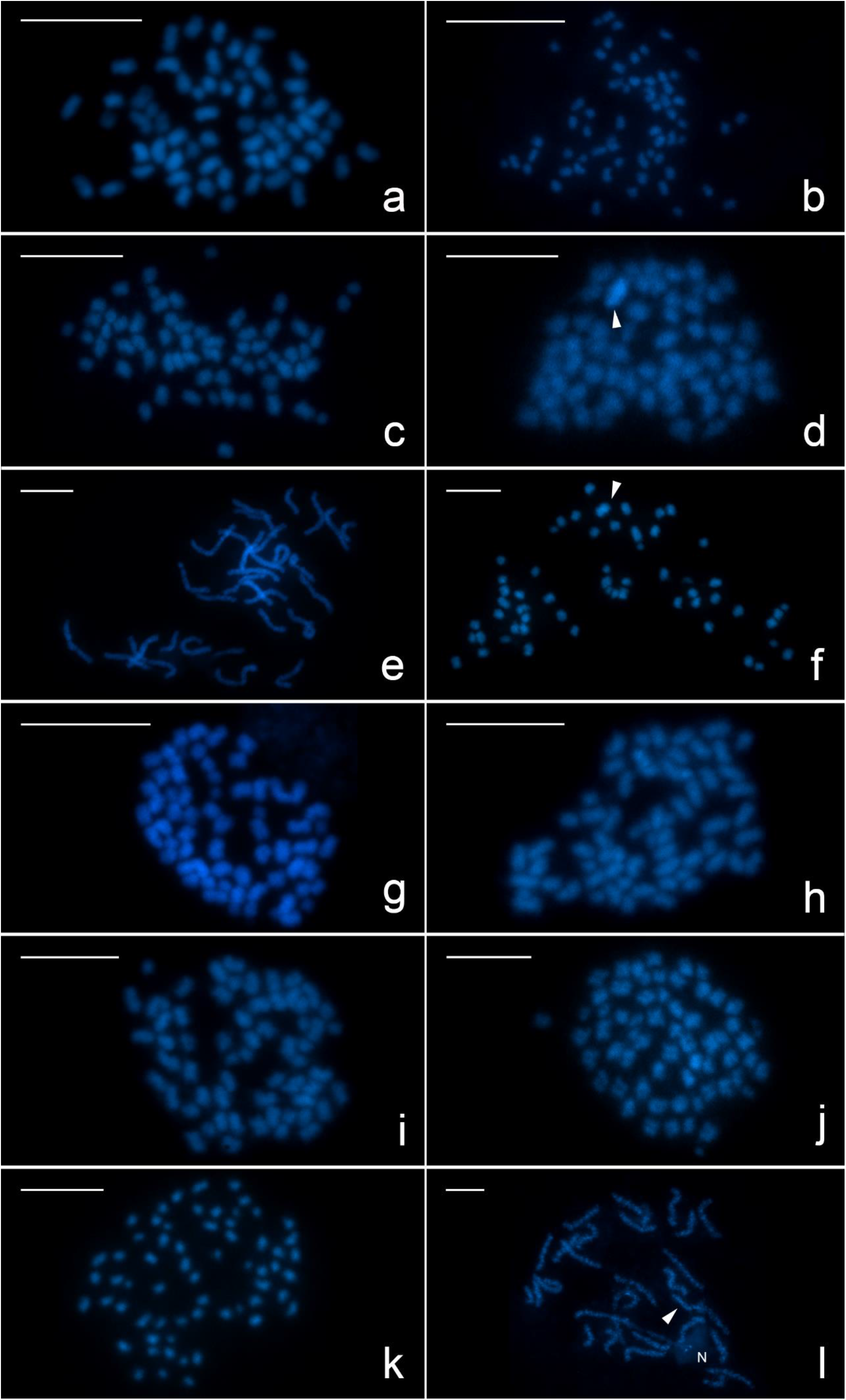
Male and female karyotypes of five *Yponomeuta* species and their outgroup *Teinoptila gutella*. Chromosomes are counterstained by DAPI (blue). a - male mitotic nuclei of *Teinoptila gutella* (2n=62), b - female mitotic nuclei of *Teinoptila gutella*(2n=61), c - male mitotic nuclei of *Yponomeuta plumbella* (2n=62), d - female mitotic nuclei of *Yponomeuta plumbella* (2n=61), e - male pachytene nuclei of *Y. polysticta* (2n=62), f - female mitotic nuclei of *Y. polysticta* (2n=61), g - male mitotic nuclei of *Y. kanaiella* (2n=62), h - female mitotic nuclei of *Y. kanaiella* (2n=61), i - male mitotic nuclei of *Y. orientalis* (2n=62), j - female mitotic nuclei of *Y. orientalis* (2n=61), k - male mitotic nuclei of *Y. mahalebella* (2n=62), k - female pachytene nuclei of *Y. mahalebella* (2n=61). N - nucleolus, W chromosome is indicated by arrowhead. Scale = 10 µm.

**Figure 2:**
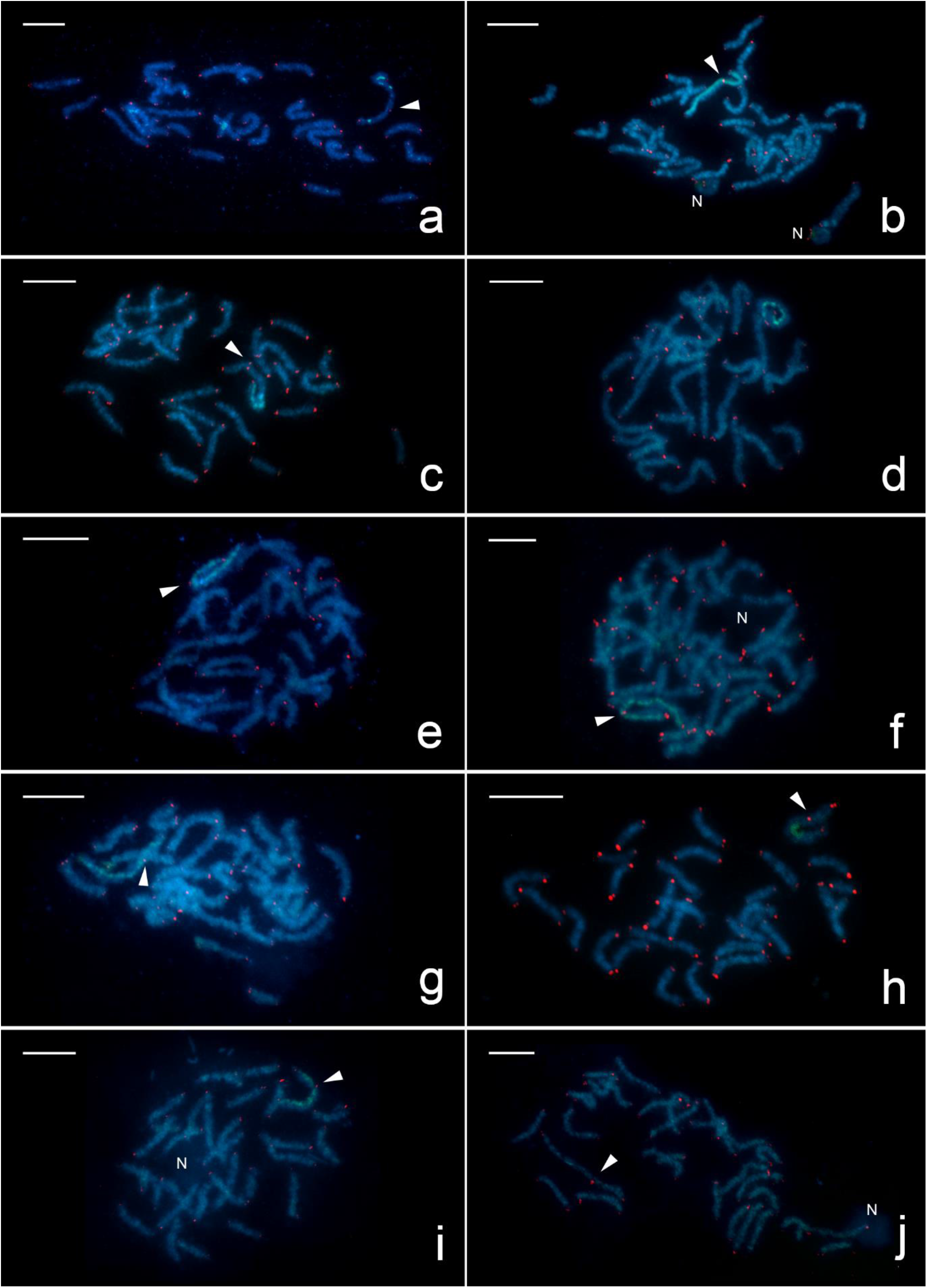
Sex chromosome trivalent (Z1Z2W) detected by GISH with female gDNA probe (green) and telomeric probe (red) on female pachytene nuclei (blue) of nine *Yponomeuta* spp. and their outgroup *Teinoptila gutella*. Telomeric signals of Z1 and Z2 chromosomes within the trivalent are marked by arrowheads. a - *Teinoptila gutella, b* - *Yponomeuta plumbella*, c - *Y. polysticta*, d - *Y. tokyonella*, e- *Y. kanaiella*, f - *Y. evonymella*, g - *Y. orientalis*, h – *Y. cagnagella*, i - *Y. padella*, j - *Y. mahalebella*. N - nucleolus. Scale = 10 µm.

### GISH with telomeric probe

To determine the constitution of sex chromosomes, genomic *in situ* hybridization (GISH) using female gDNA probe combined with telomeric probe was carried out in all studied species except for

*Plutella xylostella* where the sex chromosome system has been already known to be ZW/ZZ (Dalíková et al. 2017) The multiple sex chromosome constitution, ♀Z1Z2W/♂Z1Z1Z2Z2, was observed in all studied species except for *Y. tokyonella* with ♀ZW/♂ZZ system **(Fig. 2d**). In pachytene stage during meiosis in female ovaries both Z chromosomes, Z1 and Z2 chromosomes, pair with W chromosome forming a trivalent which is the biggest element of the karyotype. In *T. gutella, Y. plumbella, Y. kanaiella, Y. evonymella, Y. orientalis, Y. padella* and *Y. mahalebella*, the female gDNA probe hybridized evenly along the whole length of W chromosome (**Fig. 2a, b, e-g, i, j**). Whereas in *Y. cagnagella* and *Y. polysticta*, part of W chromosome was showed higher signal intensity which was probably caused by presence of compact heterochromatin of the ancestral W chromosome (**Fig. 2c, h**).

### Array-CGH

To identify Z-linked orthologs in *Y. evonymella*, we carried out array-CGH (**Fig. 3**). After filtering, *Y. evonymella* log2 ratio values of male-to-female signal intensities (log2(M>F) for 4477 orthologs were obtained. The values averaged across two replicas clearly showed a bimodal distribution (**Fig. 3**). Using a cut-off value of 0.5, we identified 245 putative Z-linked orthologs. The orthologues were assigned to chromosomes assuming a conserved synteny of genes between *Y. evonymella* and *B. mori*. The identified *Y. evonymella* Z-linked orthologs were assigned to the *B. mori* Z chromosome (BmChr1) and chromosome 2 (BmChr2; **Fig. 3**).

**Figure 3:**
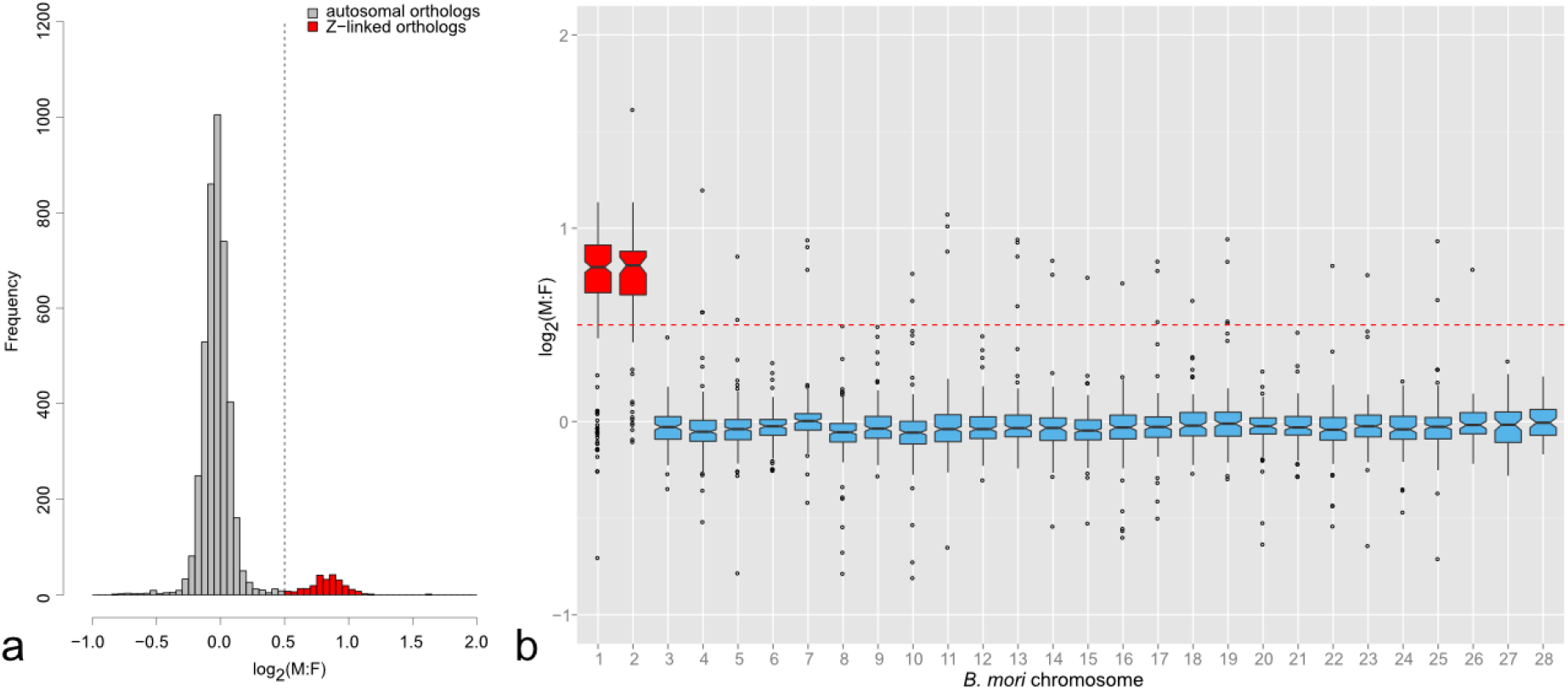
Array-CGH in *Y. evonymella*. a - Distribution of CGH log2 ratio of male-to-female signal intensities [log2(M:F)]. The peak centered at −0.05 (grey) corresponds to putative autosomal orthologs while bins with values >0.5 (dashed line) form the smaller peak comprise putative Z-linked orthologs (red). A total of 4477 orthologs are presented. b – Assignment of *Y. evonymella* orthologs to the *B. mori* chromosomes. The putative *Y. evonymella* Z-linked orthologues were assigned to chromosomes 1 (Z) and 2 of *B. mori* (red) with log2(M:F)>0.5 (dashed line). Autosomes are in blue. Boxes represent median and first and third quartiles. Whiskers extend to 1.5 * IQR from the hinges, where IQR is distance between the first and third quartiles.

### BAC-FISH

To verify results of the array-CGH analysis, we physically mapped chromosome markers to the *Y. evonymella* Z1 and Z2 chromosomes. As putative markers for the chromosome Z1, the *Y. evonymella* BAC library was screened for BACs containing single copy orthologs to genes *Henna* and *Kettin*, in *B. mori* localized on chromosome Z (BmChr1). Analogously, BACs containing orthologs of *Arp6* and Plep1 linked to chromosome 2 in *B. mori* (BmChr2) were used for the chromosome Z2. The BAC clones bearing these markers were hybridized to female meiotic nuclei of *Y. evonymella* using BAC-FISH (**Fig. 4**). BAC probes bearing Z1 marker *Henna* and Z2 markers *Arp6* and *Plep1* successfully hybridized to the sex chromosome trivalent and provided clear and discreet signals on the Z1 and Z2 chromosomes, respectively. The probe derived from BAC bearing *Kettin* hybridized to the terminal regions of all chromosomes (results not shown) which could be explained by presence of telomeric or other repetitive sequences in the BAC. The *Kettin* clone was therefore excluded from this analysis. This experiment corroborated results of array-CGH analysis and confirmed that the Z1 and Z2 chromosomes form together with the W chromosome a trivalent in *Y. evonymella*.

**Figure 4:**
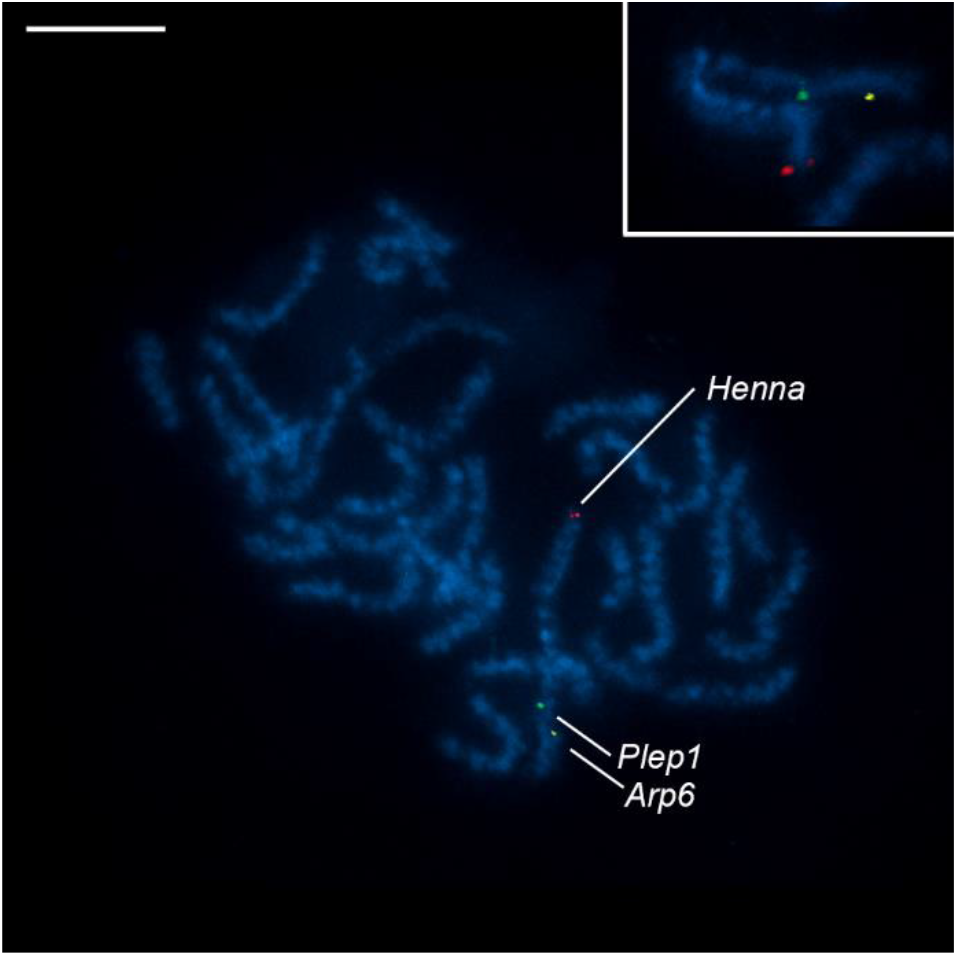
Localization of BAC clone bearing gene *Henna*, marker for Z1 chromosome (red signal), and BAC clones bearing gene *Plep 1* (green signal) and *Arp 6* (yellow signal), markers for Z2 chromosome detected on female pachytene nuclei of *Y. evonymella*. Chromosomes are counterstained by DAPI (blue). Hybridization signals are indicated by arrowhead. Scale = 10 µm.

### Quantitative PCR

To verify the hypothesis that the multiple sex chromosome system occurred in the common ancestor of the genus *Yponomeuta*, a relative gene dose of the Z1-linked genes, *Henna* and *Kettin*, and the Z2-linked genes, *Arp6* and *Plep1*, was compared by qPCR experiment between female and male gDNA of *Y. evonymella*, and *Y. plumbella* representing the early diverged *Yponomeuta* species. We also tested *Y. tokyonella* in which chromosome number was reduced to 2n=60 in both sexes. The results showed statistically significant twofold difference between males and females in all studied genes and thus proved their Z-linkage in all three *Yponomeuta* species (**Fig. 5, Suppl. Tab. S4**). The only exception was the gene *Henna* in *Y. plumbella* which showed autosomal linkage. The Z chromosome is considered conserved across Lepidoptera and the Z-linkage of the *Henna* gene has been confirmed in many species (Van’t Hof et al. 2013; Nguyen et al. 2013; Dalíková et al. 2017). It is reasonable to assume that a chromosomal rearrangement such as translocation moved the *Henna* gene to an autosome.

**Figure 5:**
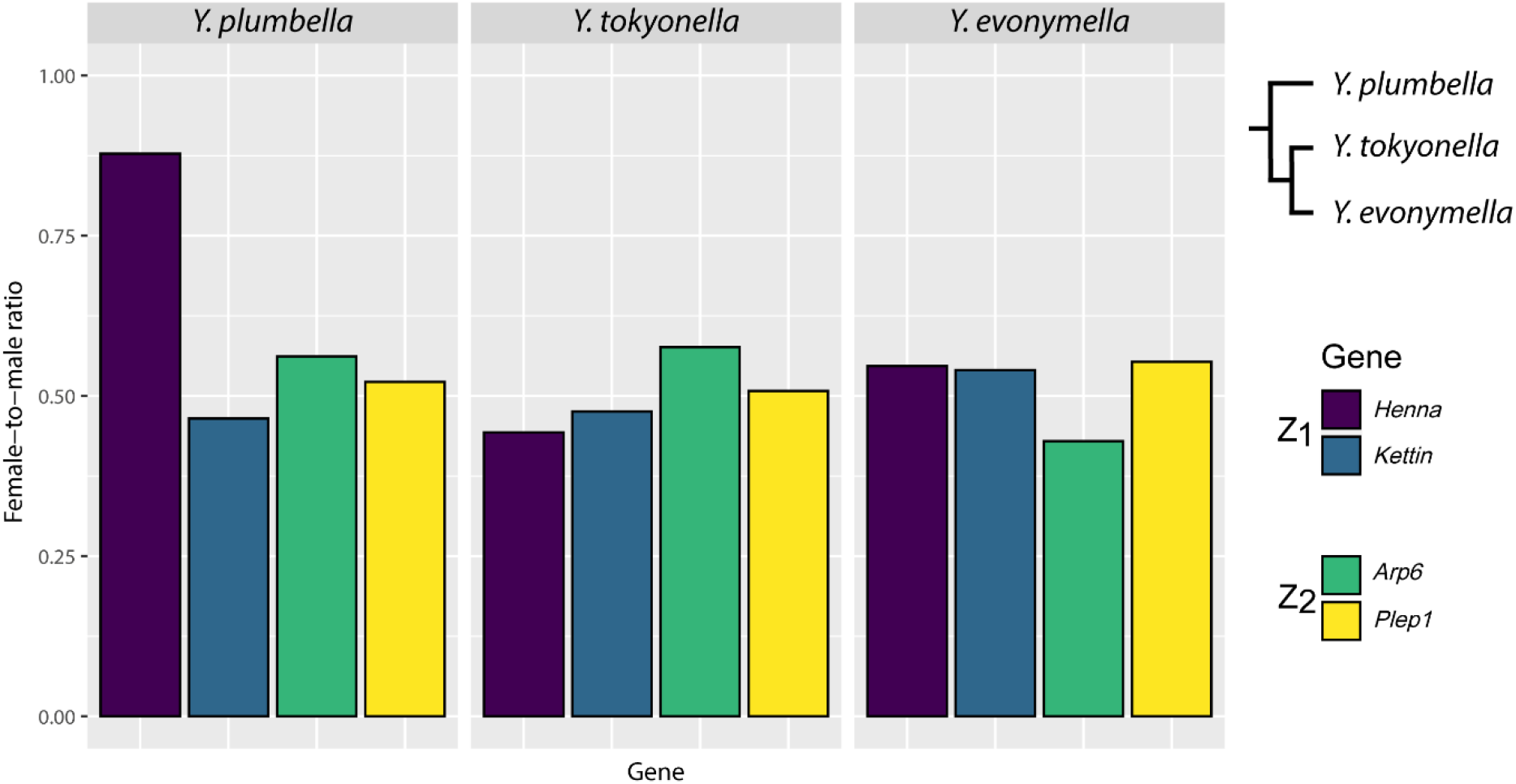
Mean female-to-male ratio values obtained by qPCR for Z1-linked genes *Henna* and *Kettin* and Z2-linked genes *Arp6* and *Plep1* using *Ace2* as the reference gene in *Y. plumbella, Y. tokyonella* and *Y. evonymella*. A value of 0.5 is expected for Z-linked genes, while for autosomal genes the expected value is 1. For summary of qPCR results, see **Supplementary Table S4**.

## Discussion

Ermine moths of the genus *Yponomeuta* represent a suitable system to study a role of changes in genome organization in a host shift. While *Yponomeuta* and *Teinoptila* spp. have an ancestral association with a single plant family, Celastraceae, the European *Yponomeuta* clade shifted to new hosts of the families Rosaceae and Salicaceae. Using the combination of GISH and FISH with telomeric probe, we detected the heterologous W chromosome in all species under study and clearly identified the multiple sex chromosome constitution ♀Z1Z2W/♂Z1Z1Z2Z2 reported by (Nilsson et al. 1988) except for *Y. tokyonella* (**Fig. 2**). Our results thus show that formation of the multiple sex chromosomes probably not played a major role in the host shift of European ermine moths as they predate the split of the *Yponomeuta* and *Teinoptila* genera. Since a representative of a sister clade of ermine moths, *P. xylostella* (Plutellidae), has an ancestral genome organization 2n=62 with the ZW sex chromosome system (Ward et al. 2021), the *Yponomeuta* multiple sex chromosomes most likely rose within the family Yponomeutidae. Analysis of additional yponomeutids is necessary to pinpoint their exact origin. Results of array-CGH confirmed by physical mapping and qPCR further revealed that the Z2 chromosome of the *Yponomeuta* multiple sex chromosome system corresponds to a synteny block homeologous to the *B. mori* chromosome 2 (BmChr2; **Fig. 3, 4, 5**).

In *Y. tokyonella*, we observed the ♀ZW/♂ZZ sex chromosome constitution, where the fusion between ancestral sex chromosomes and a pair of autosomes was completed, forming neo-W and neo-Z chromosome (♀WZ/♂ZZ; **Fig. 2d**). It was shown that autosomal fusions are not random in Lepidoptera, with small and repeat rich autosomes being repeatedly involved in distant taxa (Ahola et al. 2014). Furthermore, it was proposed that sex chromosome-autosome fusions are the first rearrangements to differentiate karyotypes from ancestral lepidopteran genome architecture (Carabajal Paladino et al. 2019). We hypothesize that sex chromosome turnover is initiated by the repeat rich W chromosome in Lepidoptera giving rise to the Z1Z2W multiple sex chromosome system such as the one observed in *Yponomeuta*. Pairing of the Z1 and Z2 chromosomes with the W chromosome increases probability of their interaction (cf. Schlecht et al. 2004), which can result in a fusion mediated by ectopic recombination and produce a conspicuously large pair of neo-sex chromosome observed in other Lepidoptera (Nguyen et al. 2013; Mongue et al. 2017; Carabajal Paladino et al. 2019).

According to the null model of sex chromosome-autosome fusions developed by Anderson et al. (2020), all chromosomes fuse with equal probability. The model showed that sex chromosome-autosome fusions make a large proportion of fusions even in absence of selection in species with small number of autosomes, while they should be rare in clades with high chromosome number such as Lepidoptera. Yet, growing number of neo-sex chromosome systems have been reported in Lepidoptera, which suggests that sex chromosome-autosome fusions are common in this female heterogametic group (Nguyen and Carabajal Paladino 2016; Carabajal Paladino et al. 2019). This is in stark contrast to analyses performed in vertebrates (Pokorná et al. 2014; Pennell et al. 2015; Sember et al. 2021), where fusions between sex chromosomes and autosomes are rare.

It was hypothesized that sex chromosome-autosome fusions are more likely to be deleterious compared to fusions between autosomes in species with achiasmatic meiosis as a result of sex chromosome differentiation process (Anderson et al. 2020). As genes cease to recombine upon a sex chromosome-autosome fusion, they start accumulating mutations. Deleterious mutations thus quickly overcome any initial fitness benefit of the fusion and prevent its fixation (Anderson et al., 2020; cf. Lenormand and Roze, 2022). The deleterious effect should be proportionate to number of genes born by the involved autosome. Thus, the high incidence of neo-sex chromosomes in Lepidoptera could be explained by their achiasmatic female meiosis and numerous small chromosomes.

The *B. mori* chromosome 2 is one of the smallest elements in the karyotype (Yoshido et al. 2005a). As mentioned above, smaller chromosomes with high repetitive DNA content are more prone to chromosome rearrangements compared to large ones which generally have less repetitive DNA (Ahola et al. 2014). A recent analysis of repetitive landscape in two *Danaus* species showed that various mobile elements are more abundant in small chromosomes (Baril and Hayward 2022). Indeed, the chromosome 2 fused with other chromosomes in *S. cynthia* (2n = 25-28) (Yoshido et al. 2011) or *Manduca sexta* (n=28) (Yasukochi et al. 2009). Notably, a similar pattern was also observed in birds. The typical avian karyotype consists of about 80 macro- and micro-chromosomes (Ellegren 2010; Zhang et al. 2014) except for some groups such as parrots (de Oliveira Furo et al. 2017) and birds of prey (de Oliveira et al. 2005; Joseph et al. 2018). Their reduced chromosome numbers are mostly caused by lineage specific fusions of microchromosomes and smaller macrochromosomes including sex chromosomes (Wilcox et al. 2019; Furo et al. 2020; Huang et al. 2022). It was hypothesized that in parrots, expansion of mobile elements could lead to chromosome rearrangements and subsequent loss of genes involved in maintaining genome stability and repair of double-strand breaks (Huang et al. 2022).

Alternatively, it was proposed that fixation of neo-sex chromosomes in lepidopteran populations could be facilitated by sexual antagonistic selection (Charlesworth and Charlesworth 1980; Smith et al. 2016; Matsumoto and Kitano 2016) or selection for linkage between largely sex-linked reproductive barriers and larval performance (Nguyen et al. 2013; Carabajal Paladino et al. 2019). As for the latter, the inspection of a gene content of the chromosome 2 in the reference genome of *B. mori* did not show any enrichment for genes involved in detoxification of plant secondary metabolites, which are crucial for performance of larvae on their host plants (cf. Yu et al., 2008; Tsubota and Shiotsuki, 2010; Ai et al., 2011; Ahn et al., 2012; Xie et al., 2012).

However, BmChr2 bears a cluster of more than 100 chorion genes, which are expressed in ovaries and encode specialized structural proteins found in the eggshell (Goldsmith and Basehoar 1978; Suetsugu et al. 2013). Notably, (Suetsugu et al. 2013) shown that 74% of genes with ovary specific expression is clustered on chromosomes 2, 10, 15, and 16 in the *B. mori* genome. While a fragment of chromosome 2 fused with a Z chromosome also in *Pieris* white butterflies (Pieridae; Hill et al., 2019; Steward et al., 2021), chromosomes 15 and 16 fused with Z chromosomes in moths of the family Tortricidae and *Danaus* spp., respectively (Nguyen et al. 2013; Mongue et al. 2017). Ovary specific expression and sex linkage are alternative ways to resolve sexual conflict (Mank 2009). Thus, we hypothesize that sex chromosome turnover in Lepidoptera could be driven by sexual antagonism, which is in agreement with theoretical predictions (Charlesworth and Charlesworth 1980; Matsumoto and Kitano 2016) and findings in a three-spined stickleback fish (Kitano et al. 2009; Dagilis et al. 2022). Further research on distribution of sexual antagonistic loci in lepidopteran genomes is needed to test this hypothesis.

## Supplementary tables

**Supplementary table S1:**
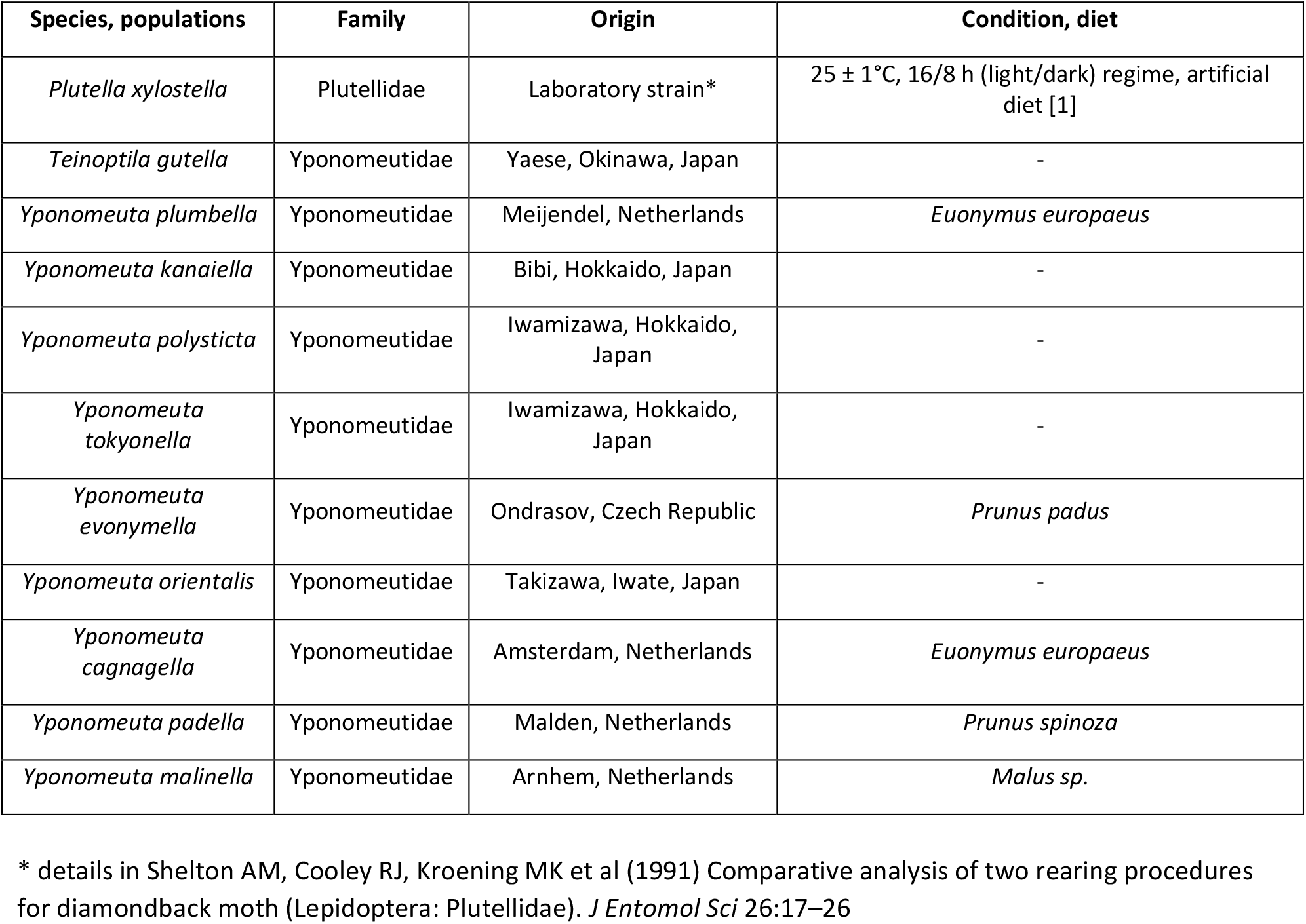
List of species examined.

**Supplementary Table S2:**
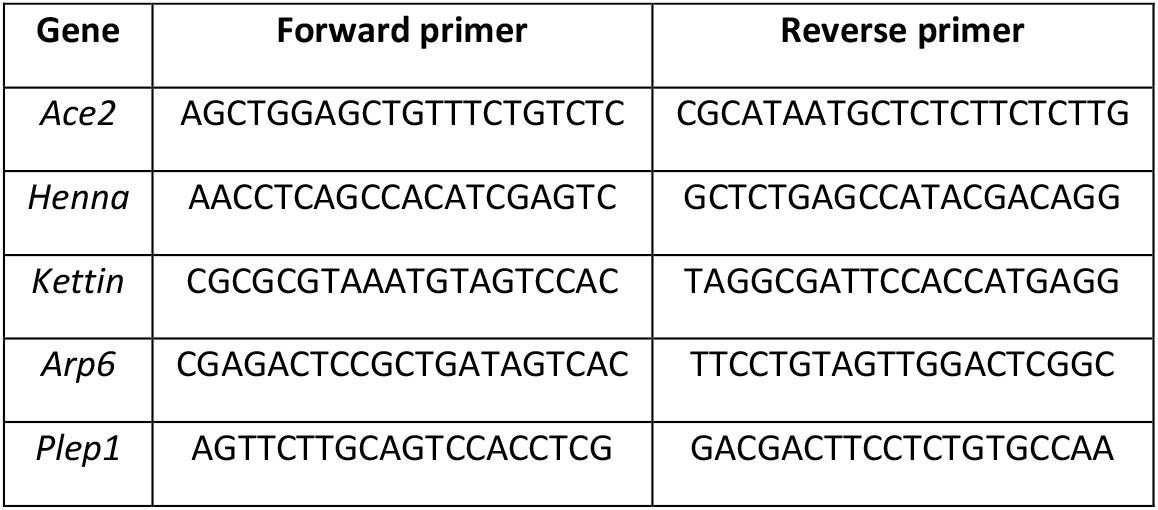
List of primers used for screening of BAC library of *Y. evonymella*.

**Supplementary Table S3:**
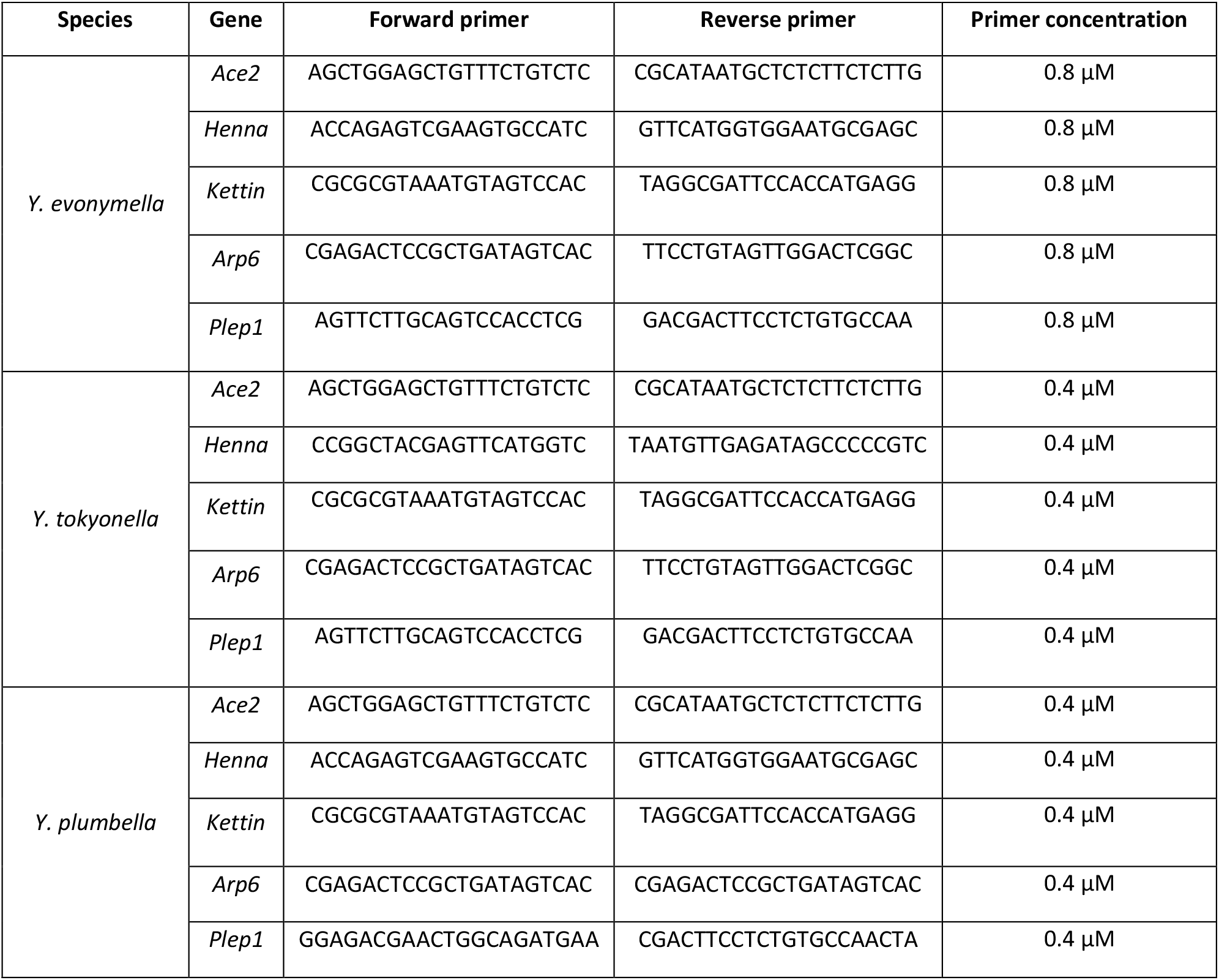
List of primers used in qPCR experiments.

**Supplementary Table S4:** Summary of qPCR results. Target gene to reference ratio (*R*) was determined in three biological samples (I-III) in males (M) and females (F) using the reaction efficiencies EReference and ETarget. The mean and its standard error (S.E.) was calculated from these three independent R values. Two null hypotheses were tested by unpaired two-tailed t-test for unequal variances. In the autosomal hypothesis (A) we tested female-to-male R ratio 1:1, whereas in the Z-linkage hypothesis (Z) the tested female-to-male ratio was 1:2. P-value lower than 0.05 means significant difference from tested ratio.

| Species | Target gene | Sex | Target gene to reference ratio ($R$) | | | | | | P value of t-test | |
| --- | --- | --- | --- | --- | --- | --- | --- | --- | --- | --- |
| | | | sample I | sample II | sample III | $E_{\text{Reference}}$ | $E_{\text{Target}}$ | mean ± S.D. | A | Z |
| *Yponomeuta plumbella* | *Plep1* | F | 0,182 | 0,175 | 0,181 | 0,877 | 0,905 | 0,179±0,004 | 0,0006 | 0,406 |
| | | M | 0,375 | 0,333 | 0,321 | | | 0,343±0,028 | | |
| | *Arp6* | F | 0,075 | 0,086 | 0,083 | 0,877 | 1,013 | 0,082±0,006 | 0,0004 | 0,1005 |
| | | M | 0,138 | 0,155 | 0,143 | | | 0,146±0,009 | | |
| | *Kettin* | F | 0,178 | 0,139 | 0,179 | 0,877 | 0,922 | 0,165±0,022 | 0,0004 | 0,441 |
| | | M | 0,331 | 0,372 | 0,362 | | | 0,355±0,022 | | |
| | *Henna* | F | 0,075 | 0,077 | 0,065 | 0,877 | 0,91 | 0,072±0,007 | 0,121 | 0,001 |
| | | M | 0,083 | 0,077 | 0,087 | | | 0,082±0,005 | | |
| *Yponomeuta tokyonella* | *Plep1* | F | 0,335 | 0,383 | 0,348 | 1,015 | 1,008 | 0,355±0,025 | 0,009 | 0,888 |
| | | M | 0,739 | 0,559 | 0,798 | | | 0,699±0,125 | | |
| | *Arp6* | F | 0,699 | 0,565 | 0,918 | 1,015 | 0,998 | 0,727±0,178 | 0,008 | 0,411 |
| | | M | 1,199 | 1,249 | 1,337 | | | 1,262±0,07 | | |
| | *Kettin* | F | 0,468 | 0,541 | 0,541 | 1,015 | 0,951 | 0,516±0,042 | 0,023 | 0,764 |
| | | M | 0,873 | 0,993 | 1,39 | | | 1,085±0,271 | | |
| | *Henna* | F | 0,455 | 0,396 | 0,312 | 1,015 | 0,999 | 0,388±0,072 | 0,0004 | 0,299 |
| | | M | 0,855 | 0,902 | 0,871 | | | 0,876±0,024 | | |
| *Yponomeuta evonymella* | *Plep1* | F | 0,245 | 0,223 | 0,217 | 0,89 | 0,96 | 0,228±0,015 | 0,002 | 0,224 |
| | | M | 0,364 | 0,45 | 0,423 | | | 0,412±0,044 | | |
| | *Arp6* | F | 0,554 | 0,568 | 0,511 | 0,89 | 0,903 | 0,545±0,029 | 0,003 | 0,196 |
| | | M | 1,439 | 1,06 | 1,31 | | | 1,269±0,193 | | |
| | *Kettin* | F | 0,13 | 0,114 | 0,158 | 0,89 | 0,996 | 0,134±0,022 | 0,012 | 0,598 |
| | | M | 0,205 | 0,284 | 0,255 | | | 0,248±0,039 | | |
| | *Henna* | F | 4,179 | 3,526 | 4,046 | 0,89 | 0,744 | 3,917±0,345 | 0,002 | 0,313 |
| | | M | 7,646 | 6,321 | 7,525 | | | 7,164±0,732 | | |

